# A visual representation of the hand in the resting somatomotor regions of the human brain

**DOI:** 10.1101/2022.04.05.486995

**Authors:** Yara El Rassi, Giacomo Handjaras, Andrea Leo, Paolo Papale, Maurizio Corbetta, Emiliano Ricciardi, Viviana Betti

## Abstract

Hands are *regularly* in sight in everyday life. This visibility affects motor control, perception, and attention, as visual information is integrated into an internal model of sensorimotor control. Spontaneous brain activity, i.e., ongoing activity in the absence of an active task (rest), is correlated among somatomotor regions that are jointly activated during motor tasks^1^. Moreover, recent studies suggest that spontaneous activity patterns do not only replay at rest task activation patterns, but also maintain a model of the statistical regularities (*priors*) of the body and environment, which may be used to predict upcoming behavior^2–4^. Here we test whether spontaneous activity in the human somatomotor cortex is modulated by visual stimuli that display hands vs. non-hand stimuli, and by the use/action they represent. We analyzed activity with fMRI and multivariate pattern analysis to examine the similarity between spontaneous (rest) activity patterns and task-evoked patterns to the presentation of natural hands, robot hands, gloves, or control stimuli (food). In the left somatomotor cortex we observed a stronger (multi-voxel) spatial correlation between resting-state activity and natural hand picture patterns, as compared to other stimuli. A trend analysis showed that task-rest pattern similarity was influenced by inferred visual and motor attributes (i.e., correlation for hand>robot>glove>food). We did not observe any task-rest similarity in the visual cortex. We conclude that somatomotor brain regions code at rest for visual representations of hand stimuli and their inferred use.

## RESULTS AND DISCUSSION

In this study, we examined whether human somatomotor cortex codes at rest for a visual representation of the hand and its inferred use. Using functional magnetic resonance imaging (fMRI), we acquired an eight-minute resting-state scan in which observers kept their eyes on a fixation point without any explicit cognitive or motor task (Figure 1A). We then presented observers with pictures of four categories of stimuli. i.e., natural hands, robot hands, gloves, and control stimuli (i.e., food) (Figure 1B-C). We were interested in testing two main predictions. First, we predicted that the somatomotor cortex would represent at rest more commonly hand vs. non-hand stimuli. This is given by the nearly continuous vision of our hands in everyday life, as compared to hand-like stimuli like robot hands or gloves. These are stimuli sharing visual attributes with hands but are much less common. Second, inferred action/use may also modulate spontaneous activity in the somatomotor cortex. Robot hands perform similar action to hands, while gloves have no autonomous motor attributes, despite their relative visual similarity. To test the association between resting-state spontaneous activity and task-evoked responses, we selected three separate regions of interest (ROIs): left and right somatomotor areas (i.e., precentral and postcentral gyri), identified with a three-minute finger tapping scan (Figure 1D), and bilateral early visual areas (V1-V2-V3), selected using a functional atlas of the visual cortex.

**Figure 1.**
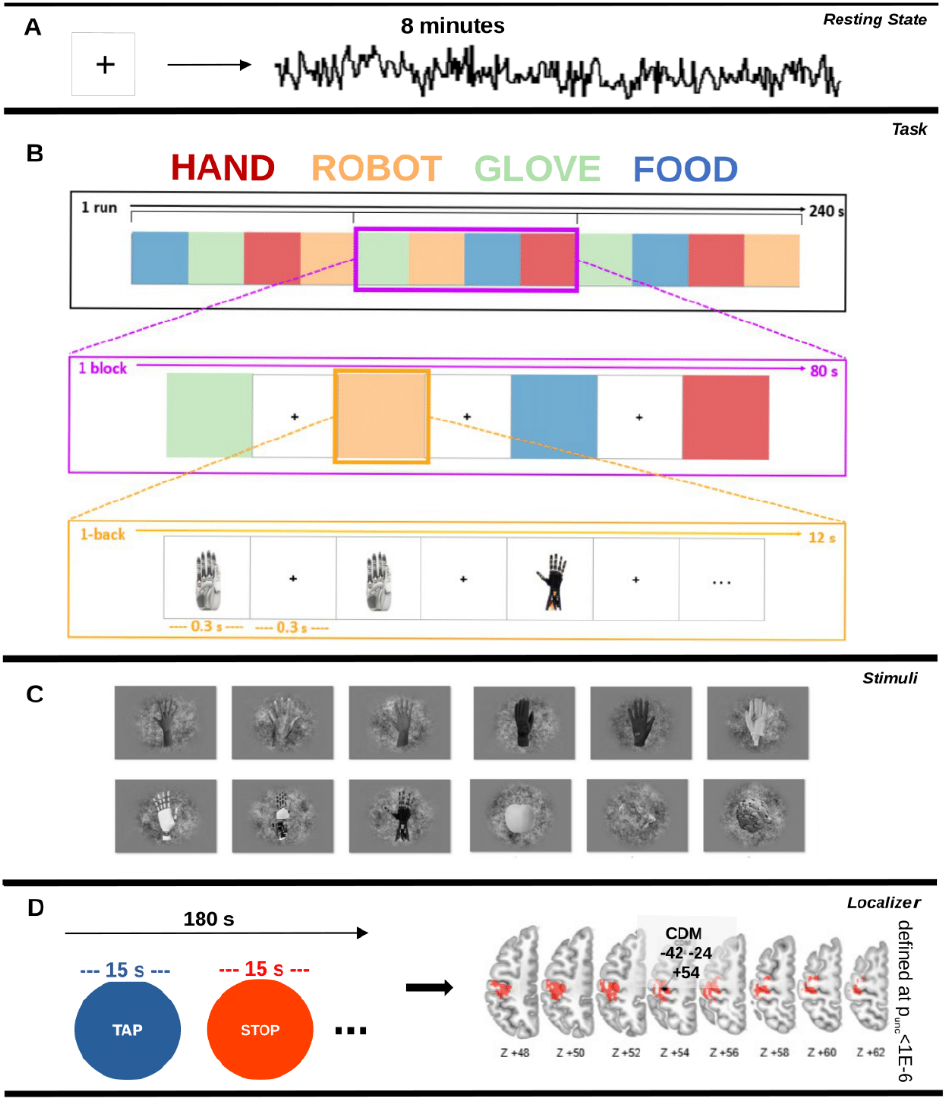
Experimental design and visual stimuli. The experiment consisted of 3 phases: i) pre-task resting-state scan (A), ii) 1-back rapid block design with four categories (B), and iii) finger tapping localizer scan (D). Visual stimuli are represented in (C).

The multi-voxel patterns of task-evoked activity of the four stimulus categories (hand, robot, glove, food) were extracted in each subject and ROI. We then correlated these task-related patterns with those extracted from each time point of the resting-state. This procedure ultimately generated a distribution of correlation coefficients for each stimulus category, ROI, and subject. We used the approach of^5^ who took the upper 90% (U90) of the distribution of correlation values to measure task-rest multi-voxel pattern similarity (Figure 2).

**Figure 2.**
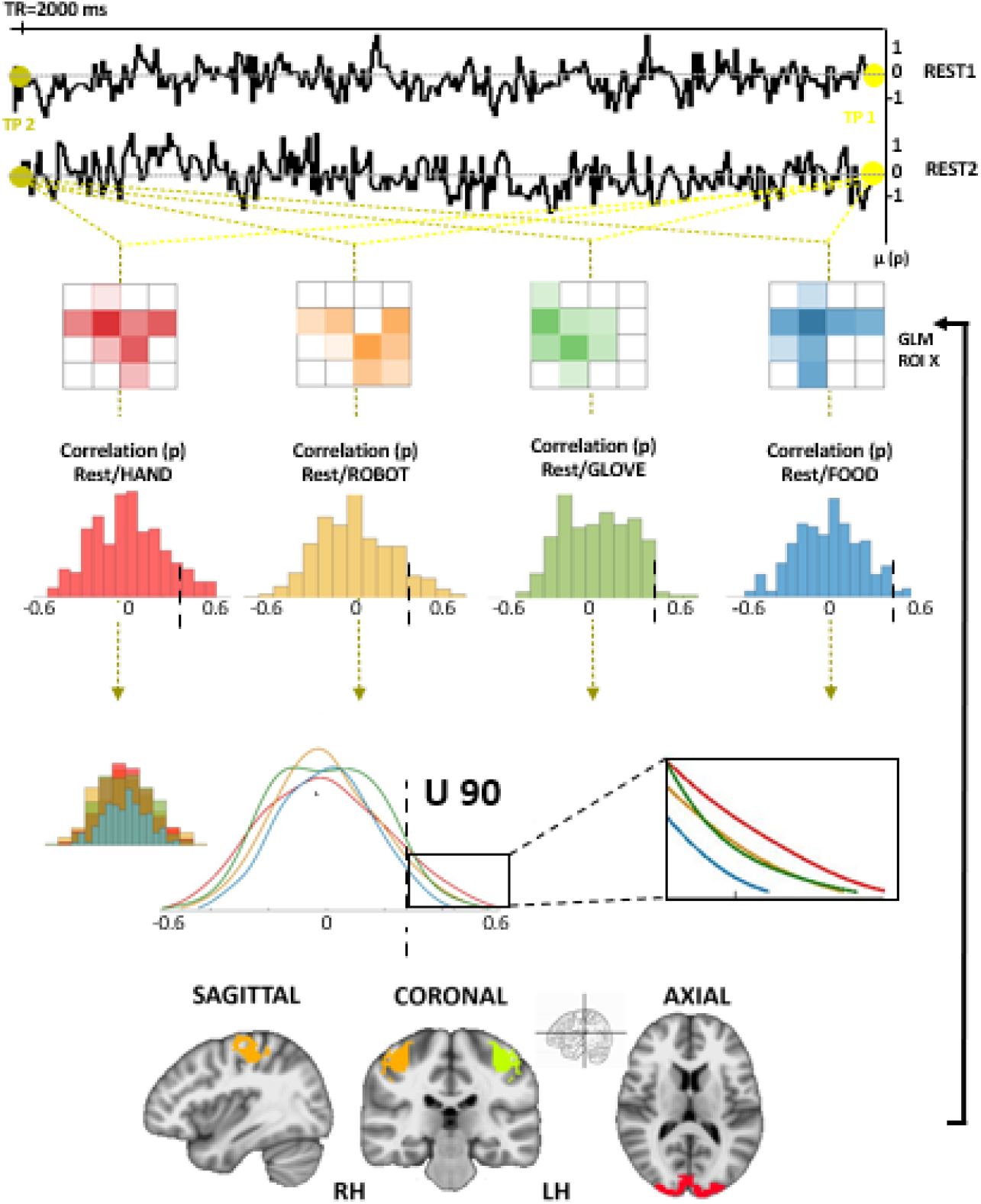
U90 analysis. We extracted the activity patterns elicited by the four stimulus categories in each subject and ROI and correlated them with those extracted from each time point of the resting state. We considered the upper 90% (U90) from the distribution of correlation coefficients for each stimulus category, ROI, and subject to measure task/rest congruency for the ANOVA. Each subject then had 4 U90 measures.

Finally, we performed a 1-way ANOVA comparing the U90 values for each category across subjects in each region of interest (left, right somatomotor, bilateral visual). In left somatomotor cortex, hand stimuli yielded the strongest rest-task similarity as compared to non-hand stimuli or objects (F(3,54)=3.469, p=.022, η=.162) (Figure 3A).

**Figure 3.**
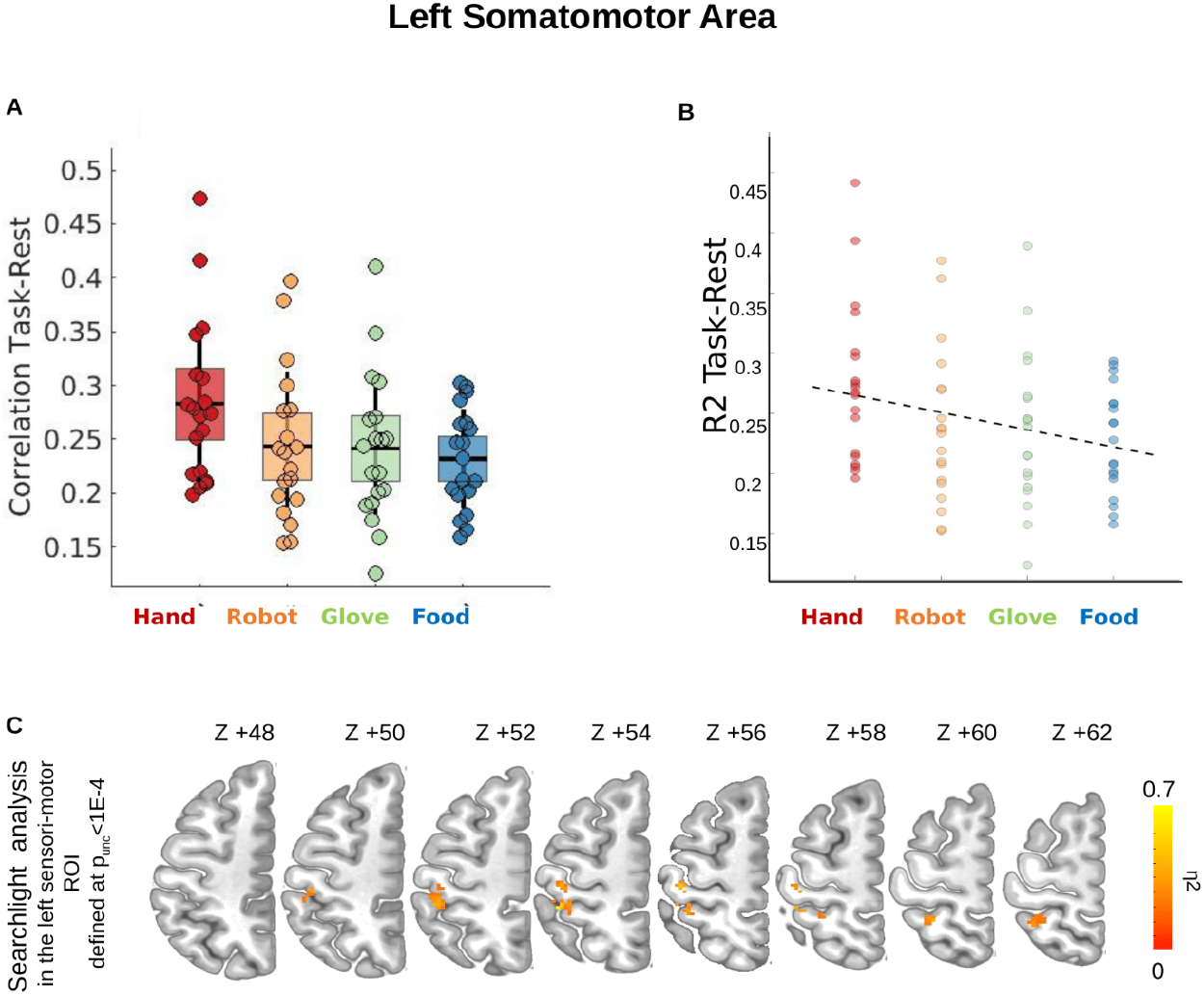
Results in the left somatomotor area. (A) ANOVA: The x-axis shows the four categories, and the y-axis shows the U90 correlation of task-rest. Each dot represents a subject. The ANOVA shows a significant main effect of visual categories (F(3,54)=3.469, p=.022, ETA=.162). (B) the graph shows a significant linear trend (hand>robot>glove>food) (F(1,18)=9.055, p=.008, ETA=.335). The x-axis shows the 4 categories, and the y-axis shows the R2 of the task-rest correlation. Each dot represents a subject. (C) The lower part shows a searchlight analysis shows the task-rest association critically depends on the postcentral gyrus activity.

The primary somatomotor cortex (S1-M1) encodes both the sensory and motor properties of body movements in a fine-grained somatotopy^6,7^. The representation structure is highly stable at the level of single fingers in S1^8,9^ and coordinated finger movements in M1^10,11^. In these early regions, the activation patterns reflect the statistics of natural hand usage^8,10^. In the case of hand, use and visibility often co-occur, with beneficial effects on behavioral performance: hand visibility improves the accuracy of volitional movements^12^, reduces the perception of pain^13^, increases the tactile perception^14^, and allows motor-visual regularities that compute the sense of agency^15^.

At a behavioral level, we commonly rely on an intrinsic model of body structure to mediate a sense of position^16,17^. Furthermore, the brain employs a standard posture or a Bayesian *prior* for guiding body-space perception and action^18^. This is interesting in light of the idea that spontaneous activity maintains statistical regularities (*priors*) to anticipate and even predict environmental demands ^2^. This hypothesis has been tested using natural visual stimuli and common cognitive tasks^5,19–22^. More specifically, during offline periods, the brain forms generic *priors* or low-dimensional representations, as categories or synergies, rather than individual instances or movements, that summarize the relative abundance of visual stimuli, objects, or motor patterns in the natural environment. Interestingly, this reduced subspace of summary representations is formed along a hierarchy that has the sensorimotor cortices at the lower level ^4^.

Consistently, our results show that at rest the somatomotor regions maintain a multivoxel pattern of activity that resembles that evoked by the presentation of the natural hands compared to control stimuli (e.g., food). Therefore, the somatomotor regions acting as a central node of processing of afferent and efferent inputs, fundamental for the active tactile feedback and proprioception, may retain representations (e.g., the body form) during the offline periods, instrumental to the interaction with the environment. We often rely on the physical properties of our body (especially the hand) to grasp and manipulate objects and the co-occurrence of sight and use contribute to generate priors tied to the actual experience.

According to the idea that things occurring nearly coincidentally in time are represented together in the cortex ^23^, i.e., cutaneous, proprioceptive, and visual signals, the co-occurrence statistics of usage and visibility may explain the responsiveness of the somatomotor regions to visually cued hands. Despite diverse spatial and temporal resolutions, previous findings in humans have demonstrated the existence of preferred tuning of single neurons to visually cued non-grasp-related hand shapes in the posterior parietal cortex ^24^. In monkeys, a substantial number of neurons in the arm/hand region of the postcentral gyrus is activated by both somatosensory and visual stimulation ^25^. Here, for the first time, we found that the rest-task similarity in the somatomotor cortex is driven by the hand form, and we can access it through a visually cued paradigm without explicit motor processing.

This aligns with the multimodal role of M1/S1 that embodies different body/motor-related representations, including the one mediated by hands. Linguistic studies show correlates of action words in the somatotopic activation of the motor and premotor cortex (e.g., ^26,27^.) Similarly, embodied cognition theories suggest that understanding action verbs is reliant on the involvement of action-related areas; this representation is found to be body-specific. For example, right- and left-handers perform actions differently and use different brain regions for semantic representation ^28^.

In summary, the stability of the spontaneous activity suggests that this set of neural signals is a possible candidate to preserve long-term models and priors of common behaviors and natural stimuli^3,4^. These prior representations are the result of statistical learning mechanisms that store the co-occurrence statistics of hand visibility and usage, instrumental to the exploration and manipulation of the surroundings.

### An interplay of visual and motor attributes modulates the match between resting and task-related activity

In the left somatosensory area, we followed up with a trend analysis for exploring post hoc the ANOVA interactions and testing the differences among stimulus categories (hand>robot>glove>food). Here, the aim is to better understand the multimodal activity highlighted by the visually induced effect. This investigation revealed a significant trend for decreased task-rest similarity in the multivoxel spatial pattern going from natural hand to hand-shaped objects (e.g., glove) (F(1,18)=9.055, p=.008, η=.335) (Figure 3B). Finally, a searchlight analysis (radius=6 mm) in a larger extent of the left somatomotor cortex (p<0.0001, uncorrected) was performed to better highlight the subregion of the postcentral and precentral gyri involved in the association between task and rest activity. Interestingly, this region falls in the post-central gyrus in correspondence with the hand notch (Figure 3C). Instead, in the right somatomotor area, the 1-way ANOVA showed no significant main effect of conditions (F_(3,54)_= .641, *p*=.592, η=.034), indicating that the effect was left-lateralized (Figure 4A). Similarly, in the bilateral early visual areas, the 1-way ANOVA showed no significant main effect of conditions (F_(3,54)_=1.423, *p*=.246, η=.073). As expected, there were no differences in correlation in the non-hand-preferred early visual areas (Figure 4B) that extract low- to mid-level visual features controlled for in our stimuli ^29,30^. For example, similarly, Kim and coworkers^5^ show that the visual cortex codes intrinsic representations that are specific for the stimuli being coded in the area (for example, FFA for face stimuli).

**Figure 4.**
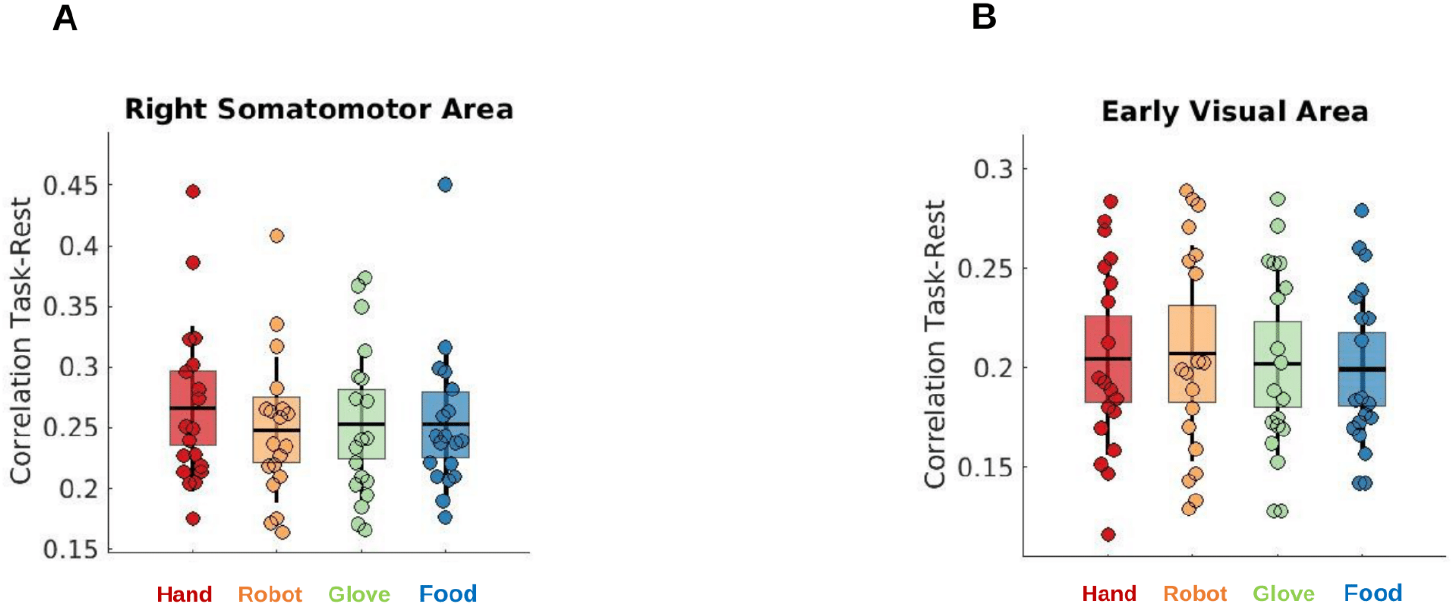
Results in right somatomotor and early visual regions. ANOVA: The x-axis shows the 4 categories, and the y-axis shows the U90 correlation of task-rest. Each dot represents a subject. In both the right somatomotor (A) and early visual areas (A), the ANOVA showed no significant main effect of conditions (p<0.05).

From birth, humans learn to use their hands in a more refined and precise fashion to interact with external objects. Spontaneous activity has been hypothesized to reflect recapitulation of previous experiences or expectations of highly probable sensory events. More precisely, the ongoing activity could be related to the statistics of habitual cortical activations during real life, both in humans^5,21,31^ and animals ^32,33^. For example,^22^ a higher similarity between motor and spontaneous patterns have been shown for natural hand sequences than for novel sequences. Based on these findings, our results can be interpreted as evidence that rest-task similarity reflects natural stimuli, or more specifically, hand-like objects as compared to artificial ones. More recent studies have found that this effect is higher in stimuli-selective regions^5^.

Beyond being natural, hands have sensory and motor attributes. From a visual point of view, gloves and robotic hands share with the hands both size and shape, but they are non-living items with either synthetic motor properties such as the robot, or no independent motor attributes such as the glove. While the visual attributes may explain the similarity between rest and task activity induced by the natural hand compared to control objects (i.e., food) (Figure 3A), the inferred action/used can bias such similarities along a continuum where hands are higher, as tied to natural movements, than robotic hands performing similarly, yet unnatural, and gloves, without autonomous motor attributes (Figure 3B). The interplay of these factors offers an interpretation of the rankings obtained: on top of the continuum, the natural hand, most abundant environmentally, has necessary visual features and motor attributes, then the robot hand though not as environmentally abundant, has the same visual features and synthetic motor attributes, the glove retains only the visual features but cannot act on its own, and finally the food objects neither have the same visual nor motor attributes.

Studies in the visual cortex demonstrated that the long-term natural experience shapes the response profile. The high-level cortical representations of these regions capture the statistics with which visual stimuli occur ^34,35^. Furthermore, animal studies demonstrated that when visual stimuli are natural scenes, the reliability of visual neurons’ response increases and persists in the subsequent spontaneous activity; these effects are not observed with the stimulation with the noise of flashed bar stimuli^32^. In the ferret’s visual cortex, the tuning function of neurons “learns” the statistics of natural but not artificial stimuli as the animal grows ^33^. Here, for the first time, we found evidence of visual representations encoded in non-visual regions at rest, but regions still specific to our hand stimuli (hand notch area). Thus, we believe that the cumulative impact of the statistics with which natural stimuli occur during the development and the experience shape the ongoing activity of the whole brain, not limited to the visual cortex. Our results are confirmed and opposed by Stringer and coworkers ^36^ that show representations of natural motor sequences at rest in the mouse visual brain but also across the forebrain. Similar to our results, they confirm that natural sequences are coded in resting state and shape the activity of the whole brain, but conversely, they find motor sequence representations in the visual system. The discrepancy could be explained by the fact that our stimuli did not represent any actions (i.e., strictly open hands), and the fact that spontaneous activity in mouse is recorded differently than in humans (mouse running in darkness vs humans staring at a cross with eyes open).

### The representation of the hand in the sensorimotor area is left-lateralized

Our study shows that the multi-voxel activity of hands in the sensorimotor area is most represented in resting-state activity; this effect was lateralized to the left, not the right, sensorimotor area (Figure 4B). From a theoretical point of view, the lateralization result is well aligned with the existing literature: compared to other body parts and objects, static pictures of hands and tools have overlapping activations in the LOTC that are then selectively connected to the left intraparietal sulcus and left premotor cortex ^37–39^. Moreover, a body of literature shows bilateral motor cortical activations are produced with the left non-dominant hand. In contrast, movements with the dominant right hand induce only contralateral (left) activations ^40^. Precisely, visuospatial orientation attention, measured with eye movements, activates a network of premotor and parietal areas in the right hemisphere, while motor attention and selection, measured as the attention needed to redirect a hand movement, activate the left hemisphere ^41,42^. TMS and lesion studies further support this left lateralization, describing motor selection, motor attention, or motor learning ^40,42^. The effector independent activation in the left hemisphere is also found in kinesthetic motor imagery that activates common circuits for motion in the premotor, posterior parietal, and cerebellar regions ^41^. More recently, Karolis and colleagues ^43^, using fMRI, built a lateralization functional taxonomy along four axes representing symbolic communication, perception/action, emotion, and decision-making. Along the action/perception axis, the categories movement, finger tapping, motor observation, and touch were all found to activate the sensorimotor areas of the left hemisphere selectively. Interestingly, all these categories had the term hand or finger as the principal components with the highest loadings.

In summary, we provide the first evidence that the ongoing activity in the left sensorimotor regions maintains a long-term representation of the hand shape in the absence of any motor task or sensory stimulation. Furthermore, this result may lend support to the representation of visually-related information in M1/S1 enforcing a multimodal role in these areas.

## CONCLUSION

The sensory and perceptual analysis is not only dependent on external stimuli but also on the coding of expected features in the surroundings simultaneous to the flow of information ^44^. Building models of the environment create priors that are stable and common across individuals yet malleable with experience and age^3^. If motor-sensory interactions are entrained throughout development into spontaneous cortical oscillations, then our internal model must have a reservoir of natural behaviors. Our experiment shows that spontaneous activity representing the internal model, despite its apparently noisy structure, reliably encodes the visuospatial topography of the natural human hand in sensorimotor areas suggesting that the human hand represents a *prior* for the effective motor interaction with the external environment to allow exploration, learning, and adaptation. In line with the malleability of the cerebral cortex in response to behavior and other input manipulations, to our knowledge, this is the first experiment to show that visually-conveyed representation of hands in resting-state activity in frontoparietal sensorimotor areas by looking at the relationship between evoked and spontaneous activity. By measuring coherence between evoked activity and resting state activity, we shed light on a multimodal role of the sensorimotor areas.

## Acknowledgments

This project has received funding from the European Research Council (ERC) under the European Union’s Horizon 2020 research and innovation programme (grant agreement No 759651) to VB. We thank Stefania Bracci for her comments to the early version of this manuscript.

## Author Contributions

Y.E.R., A.L., P.P., E.R. and V.B. developed the study design. Y.E.R. programmed the experimentation scripts. Y.E.R, G.H., and A.L. developed the pre-processing and analysis script. Y.E.R., and A.L., tested the participants. Y.E.R. and V.B drafted the manuscript. E.R. and M.C provided critical revisions. All authors revised the manuscript and approved its final version.

## Methods

### Subjects

Twenty healthy individuals (10 females) were enrolled in the study (mean age ± SD 29 ± 2.59, range 24-34). All subjects underwent medical interviews and examinations to rule out the history or presence of any disorders that could affect brain function and development. Participants were provided with a detailed description of all the experimental procedures and were required to sign a written informed consent. The study was conducted under a protocol approved by the local ethical committee (protocol n. 1485/2017) according to the Declaration of Helsinki ^45^. All subjects were right-handed, following the Edinburgh Handedness Inventory. We discarded one subject from subsequent analyses because of excessive movement artifacts in fMRI data, leading to a final sample of nineteen subjects.

### Design and Stimuli

The paradigm was divided into three parts (Figure 1). The first part included an eight-minute resting-state scan (pre-task scan). During the resting-state scan, subjects fixated a red cross (24 × 24 degrees) at the center of the screen (VisuaStimDigital dual-display goggles, 32 × 24 degrees of visual angles, Resonance Technology Inc.), without performing any cognitive or motor tasks.In the second part, we presented subjects with pictures of four categories of objects, natural hands, robot hands and control stimuli (i.e., food). All stimuli had the same vertical orientation. Left and right hands presented from the front and from the back were randomly selected and presented. We used six different examples for each object category. We presented subjects with five four-minute runs: in each run, subjects attended twelve randomized blocks (three categories, four blocks each). Each block lasted 12 seconds, followed by 8 seconds of fixation, and included 20 randomized repetitions of the six category stimuli, each presented for 0.3 seconds and followed by fixation for 0.3 seconds. Participants were instructed to perform a covert working memory one-back task as in the first task. Each run began and ended with 20 seconds of rest to acquire baseline fMRI activity. For the stimulus set, we used pictures of items pertaining to the four categories, which were converted to grayscale and matched for global luminance and root-mean-squared (RMS) contrast to control for low-level visual biases. The object pictures were centered and embedded in a circular pink-noise display with a fixed circumference (16 degrees) blending into a grey background.

For the third part, subjects performed a three-minute finger tapping localizer scan. We instructed them to tap their thumb on every other finger of their right-dominant hand sequentially and at their own pace (blocks of 15 seconds of activity followed by 15 seconds of rest).

All the visual stimuli were presented using MR-compatible display goggles (VisuaStimDigital, Resonance Technology Inc.) covering 32 × 24 degrees of visual angle, and a PC running MATLAB (MathWorks Inc, Natick, MA, USA) and the Psychophysics Toolbox version 3 ^46^.

### MRI Data Acquisition

We used a Philips 3T Ingenia scanner with a 32-channel phased-array coil. We used a gradient recall echo-planar (GRE-EPI) sequence with TR/TE = 2,000/30ms, FA =75°, FOV = 256 mm, acquisition matrix = 84 × 82, reconstruction matrix = 128 × 128, acquisition voxel size = 3 × 3 × 3 mm, reconstruction voxel size = 2 × 2 × 3 mm, 38 interleaved axial slices, and 240 volumes. We also acquired three-dimensional high-resolution anatomical images of the brain using a magnetization prepared rapid gradient echo sequence (MPRAGE) with TR/TE = 7/3.2ms, FA = 9°, FOV = 224 mm, acquisition matrix = 224 × 224, acquisition and reconstruction voxel size = 1 × 1 × 1 mm, 156 sagittal slices.

### Preprocessing

We used the AFNI software package ^47^ and a standard preprocessing pipeline to preprocess fMRI data, separately for tasks (i.e., visual working memory and finger tapping) and resting-state runs. We temporally aligned the runs (3dTshift), then corrected for head motion (3dvolreg), and used the transformation matrices to compute the framewise displacement, identifying time points affected by excessive motion^48^. We then spatially smoothed the data using a Gaussian kernel and an iterative procedure up to 6 mm Full Width at Half Maximum (3dBlurToFWHM). We then normalized the runs by dividing the intensity of each voxel over its mean over the time series and applied a multiple regression analysis (3dDeconvolve) to estimate the activation patterns for each category. We aimed to show that the hands’ category would be more strongly represented in resting state in left frontoparietal hand-preferred areas. For each subject, we modeled a GLM that included the four stimulus categories (hand, robot, glove, food) for the visual working memory task and the finger movement blocks for the hand motor localizer. The output was a stimulus-evoked BOLD multi-voxel task (category) beta weight. We included head movement parameters, framewise displacement, and signal trends as nuisance variables. We performed a generalized least squares time series fit for the resting-state data to account for nuisance regressors defined above and signal autocorrelation using an autoregressive moving average model (order 1). We registered single subject results and preprocessed rest scans to MNI152 standard space ^49^ using nonlinear registration.

### ROIs

We selected three regions of interest (ROIs) to test the association between resting-state spontaneous activity and task-evoked activity. We identified the left somatomotor area after thresholding (p<10^−6^ uncorrected) the finger tapping localizer, resulting in a small ROI (~5,000 μL) encompassing the precentral and postcentral gyri, as also suggested by the overlap with the HCP atlas^50^. The center of gravity of our ROI is −42 −24 54 (MNI atlas) and is only 6.34 mm far apart from the hand knob considering the classical paper of Chouinard and colleagues^51^ (−38 −28 52) and 8.75 mm considering Yoshimura and colleagues (2017) `(−34±4, −25±3, 57±11)`. Then, we selected the right somatomotor area by left/right flipping the ROI mentioned above (3dLRflip). Lastly, we defined the bilateral early visual cortex (V1, V2, V3) using VisFAtlas^52^ as a further early sensory control region.

### Multi-voxel pattern analysis

In the main analysis, we analyzed the association between task-evoked and resting-state activity using the procedure introduced by Kim and colleagues^5^ (Figure 2). Briefly, the patterns of task-evoked activity of the four stimulus categories (hand, robot, glove, food) were extracted in each subject and ROI. We then correlated these task-evoked patterns with the ones extracted from each time point of the resting-state. This procedure ultimately generated a distribution of correlation coefficients for each stimulus category, ROI, and subject (Figure 2). As in ^5^, we then took the upper 90% (U90) of each distribution to measure task/rest congruency. In the main analysis, we performed for each ROI an ANOVA, modeling REST and TASK (4 levels, according to the four stimulus categories). Follow-up analyses included a trend analysis in each ROI to better highlight the subregions involved in the effect of interest. This trend analysis was used to test *a priori* hypotheses of differences among stimulus categories (hand>robot>glove>food) according to empirical data, an analysis suited for exploring post hoc the ANOVA interactions. Finally, we performed a searchlight analysis (radius=6 mm) in a larger extent of the left somatomotor cortex (p<0.0001, uncorrected) to better highlight the subregions of the postcentral and precentral gyri involved in the association between task and rest activity.

